# Application of post glycosylation modifying enzymes for mass spectrometry imaging of modified *N*-glycans *in situ*

**DOI:** 10.1101/2025.05.14.654078

**Authors:** Julia E. Dreifus, Léa Chuzel, Edwin E. Escobar, Grace Grimsley, Hongxia Bai, Andrew J. Hanneman, Samantha L. Fossa, Vanessa Gregory, Richard J. Martin, Richard R. Drake, Jeremy M. Foster, Christopher H. Taron

**Affiliations:** New England Biolabs, Ipswich, MA 01938, USA; Medical University of South Carolina, Charleston, SC 29425, USA; Iowa State University, Ames, IA 50011, USA

**Keywords:** Post-glycosylation modification, mass spectrometry imaging, modified *N*-glycans, biomarker screening

## Abstract

Glycans are essential components of cells and are involved in innumerable biological processes. Their structural diversity and complexity present unique analytical challenges. Glycans are comprised of various types of monosaccharides that are linked together at different positions and with varied stereochemistry. In addition, glycans are frequently decorated with a diverse set of chemical modifications, termed post-glycosylation modifications (PGMs). Characterization of PGMs is essential for a thorough understanding of glycans, however, the technical challenges and low throughput of current methodologies have limited our understanding of these modifications. Here we demonstrate a novel approach for rapid visualization of specific PGMs present in tissue *N-*glycans by applying PGM-targeting enzymes to mass spectrometry imaging (MSI). The method enables *in situ* investigation of glycans with PGMs *en masse*, identifying the sugar residue and position modified, as well as visualizing the spatial distribution of each modified *N*-glycan in tissues. As the repertoire of PGM-targeting enzymes expands, we anticipate this approach will enable a better understanding of PGM distribution within a dynamic *N*-glycome. This may yield both new biological insights and the potential for identification of novel disease biomarkers.

## INTRODUCTION

Glycans are saccharide-based polymers that are essential to countless biological and pathological processes. These encompass structural functions as well as modulation of intrinsic and extrinsic interactions, including cell signaling, receptor ligand binding, protein folding and stability, and surface adhesion^1^. Glycan functions are determined by their composition and structure, which is defined by the types of monosaccharides that constitute them, their order of arrangement, and the linkages between units.

Glycan structural complexity is further compounded by the presence or absence of chemical modifications to individual monosaccharides, termed post-glycosylation modifications (PGMs). PGMs encompass a range of diverse chemical groups, including sulfate, phosphate, acetyl, methyl, phosphorylcholine, aminoethylphosphonate, and phosphoethanolamine, and the distinct chemistry of each alters how glycans interact with surrounding molecules^2^. Numerous diseases are associated with changes in PGMs, including autoimmune disorders, some cancers, cystic fibrosis, as well as congenital disorders^2^. As a result, PGMs represents a promising avenue of study for therapeutic and diagnostic discovery. There is wide diversity in the glycan classes PGMs modify, the monosaccharides modified, and the carbon positions occupied^2–5^. Unfortunately, with current methodologies, PGMs are difficult to identify and fully characterize, limiting our understanding of PGM distribution and how PGM patterns reflect disease states.

Existing glycan analysis methods that identify both the modified monosaccharide and the carbon-position of the modification are laborious and technically challenging. Common methods used to profile and characterize glycan mixtures involve mass spectrometry (MS), including liquid chromatography coupled with tandem MS (LC-MS/MS) and matrix assisted laser desorption ionization-time of flight (MALDI-TOF). However, while the mass of a glycan may suggest the presence of a PGM, it does not indicate the sugar residue or the specific carbon it is attached to. Additional methods are then necessary to fully investigate these samples since glycans with otherwise identical structures frequently contain isomeric PGMs^6^. Tandem MS (MS/MS) and NMR are used to fully characterize PGM positioning on glycans, however these methods are technically challenging, time-consuming, and limited to analysis of a single glycan at a time^7,8^. Moreover, fragments from PGM-containing glycans often produce overlapping *m/z* signals and may undergo in-source fragmentation, complicating confident structural assignments. Additionally, differential ionization efficiency and ion suppression effects may limit the detection of certain glycan-PGM species, particularly low-abundance or multiply modified forms. As such, these methods are not readily applicable for *in situ* analyses on cell or tissue surfaces.

In recent years, the endoglycosidase PNGase F has been applied to matrix assisted laser desorption/ionization mass spectrometry imaging (MALDI-MSI), for rapid spatial analysis of *N*-glycans in sectioned tissues^9,10^. This approach provides an *in situ* snapshot of the structures and relative abundance of *N*-glycans present as well as their localization and distribution throughout the tissue^11^. When superimposed with tissue histopathology analyses, MSI can both spatially map the *N*-glycome and visualize how it may change between healthy and diseased tissues. Since PGMs can vary across cell types and tissue microenvironments, spatially resolved methods are essential for understanding their biological roles and diagnostic potential. Such a holistic view of the dynamic *N*-glycome will better inform both basic biology and enable biomarker discovery. However, MSI exploration of PGMs in the *N*-glycome is still technically complicated. For instance, recent MSI studies have observed putative sulfated *N*-glycans, however individual MS/MS was required to validate each structure and was not practical to perform for all modified *N-*glycans observed^12,13^.

To address the technical challenges with PGM identification in MSI, we utilized a variety of highly specific enzymes that target different classes of PGMs to facilitate their rapid identification *in situ*. We first show the use of an enzyme that targets phosphorylcholine *N*-glycan modifications to visualize their spatial distribution in transverse sections of a parasitic nematode using MSI. We then illustrate the use of a sulfate-specific glycosyl hydrolase and several specificities of sulfatases to map mammalian *N*-glycan sulfation using MSI. We also present identification of new sulfatases that remove sulfate from the 3-carbon or 6-carbon of terminal galactose residues, a novel specificity. Our study both validates the use of PGM-targeting enzymes to spatially visualize different PGMs using MSI and further expands the repertoire of analytical enzymes for emerging MSI workflows.

## MATERIALS AND METHODS

### Protein expression and purification

The GlcNAc-6-phosphodiesterase and GlcNAc-6-SO_4_-hexosaminidase (F10-ORF19p) were purified as previously described^3,14^. The exo-galactose-3-sulfatase (BT1636p), exo-galactose-6-sulfatase (BT3109p), endo-galactose-3-sulfatase (BT4683p), exo-galactose-3/6-sulfatase (N4-ORF16p), and exo-galactose-3/6-sulfatase (E8-ORF27p) were cloned into the vector pJS119K with a C-terminal 6xHis tag using the NEBuilder® HiFi DNA Assembly Cloning kit (NEB, USA E5520)^15^. Primers were designed using the NEBuilder® online resource (https://nebuilder.neb.com) (**Table S1**). The genes for BT1636, BT4683, and BT3109 were cloned without the DNA encoding their predicted signal peptide sequences (residues 2-21, 2-21, and 2-17, respectively). Site-directed mutagenesis was performed using the Q5 Site-Directed Mutagenesis Kit (NEB, USA E0554S) was used to convert the catalytic serine to a cysteine (S57C) in BT3109p to enable conversion to formyl glycine in *E. coli* as previously described^16^. The resulting plasmids were used to transform NEBExpress® Competent *E. coli* cells (NEB, USA C2523). Cultures were inoculated in 1 L of LB containing 50 µg/mL kanamycin and grown at 37°C with shaking at 225 rpm. Once the culture reached exponential phase (OD_600nm_ = 0.4-0.6), protein expression was induced with 0.4 mM IPTG. Induction was performed overnight at 37°C for N4-ORF16p, BT3109p, and BT4683p. Induction was performed overnight at 30°C for BT1636p and E8-ORF27p. Cells were harvested and resuspended in binding buffer (20 mM sodium phosphate, 300 mM NaCl, 20 mM imidazole, pH 7.4). Lysis was performed using a Shearjet HPL60 High Pressure Homogenizer (Maximator, Nordhausen, Germany) with three passes at 18 kPsi. Proteins were purified on a HisTrap Fast Flow column (Cytiva, USA 17525501). Protein was eluted from the column with 500 mM imidazole. The resulting proteins were buffer exchanged into an imidazole-free buffer (20 mM sodium phosphate, 300 mM NaCl, 1 mM EDTA, pH 7.4) using Zeba Spin Desalting Columns, 7k MWCO (ThermoFisher, USA 89894).

### Sample preparation for MALDI-MSI

#### Ascaris suum

Adult *A. suum* nematode specimens were sourced from pigs, washed, and frozen at -80°C. Portions of female worms (1 cm) were mounted using a drop of Neg-50 frozen section medium (Epredia, USA) and transverse sections where cut at -20°C on a Leica GM1860 cryostat (Leica Microsystems GmbH, Wetzlar, Germany) at a thickness of 12 µm. Sectioning was performed using Cryofilm type 3C(16UF) tape and the tape containing the section was affixed with SCMM-MS1 mounting medium (SECTION-LAB Co. Ltd., Yokohama Kanagawa, Japan)^17^.

#### Human serum

Human serum (Sigma, P2918) was mixed 2:1 with 100 mM sodium bicarbonate, pH 8.0 and spotted onto an amine-reactive slide (Grace Bio-labs, 246865), using a 64 well ProPlate Slide Module to guide spot placement (Grace Bio-labs, 63484-48)^18^. Spots were immobilized on the slide by immediately placing the slide in a preheated humidity chamber (37°C) and incubating at room temperature for 1 hour. The slide was then placed in a desiccator for 15 minutes to dry the spots. To desalt the samples, the spots were incubated in Carnoy’s solution (10% glacial acetic acid, 30% chloroform, 60% ethanol) for 3 minutes, three times. After rinsing with HPLC-grade water, the samples were dried in a desiccator.

For the step-wise exoglycosidase digestions, α2-3,6,8,9-neuraminidase A (NEB, P0722) was applied to the sample at 0.1 mg/mL, β1-3,4-galactosidase (NEB, P0746) was applied at 0.1 mg/mL, and the GlcNAc-6-SO_4_ hexosaminidase (F10-ORF19p) was applied at 6 µg/mL. Enzymatic reactions were incubated at 37°C for 16h in a humidity chamber^19^, and samples were rinsed with water in between reactions. Slides were then rinsed sequentially in the following solutions to remove excess enzymes and salt: 1 minute in 100% ethanol, 95% ethanol, 70% ethanol, and HPLC water; 3 minutes in 20 mM Tris pH 9.0; 1 minute in HPLC water; 3 minutes in 10 mM citraconic anhydride buffer, pH 3; 1 minute in HPLC water. The samples were then dried in a desiccator.

#### Thyroid tissue

Fresh frozen cancerous human thyroid tissue was acquired from BioIVT (New York, USA). The thyroid sample was then mounted using a drop of Neg-50 frozen section medium (Epredia) and sectioned at -16°C on a Leica GM1860 cryostat (Leica Microsystems GmbH, Wetzlar, Germany) at a thickness of 12 µm. Sections were thaw-mounted onto SuperFrost Plus adhesion microscope slides (Epredia) and placed in a desiccator to dry. One of the sections was stained with hematoxylin and eosin (HCE) histological stain to visualize tissue pathology by incubating in the following solutions: 30 s in 95% ethanol, 30 s in 70% ethanol, 30 s in HPLC water, 2 minutes in Hematoxylin, 30 s in HPLC water, 30 s in 70% ethanol, 30 s in 95% ethanol, 1 minutes in Eosin-Y solution, 30 s in 95% ethanol, 30 s in 100% ethanol, and 2.5 minutes in xylenes. High resolution images of the HCE-stained tissue were taken with a Hamamatsu NanoZoomer SQ System at 20X magnification and exported at 5X magnification (Hamamatsu). Samples were subjected to washes with ethanol and Carnoy’s solution (10% glacial acetic acid, 30% chloroform, 60% ethanol) followed by antigen retrieval in 10 mM citraconic anhydride buffer, pH 3 for 5 min in a NxGen Decloaking Chamber (Biocare Medical, USA) at 110°C as previously described^10,20^. The samples were then dried in a desiccator. A hydrophobic pen was used to draw a barrier around each thaw-mounted sample. Sulfatase reactions were performed with 0.1 mg/mL enzyme in 50 mM Tris, pH 8.0 for 16h at 37°C with 5 mM CaCl_2_ or 5 mM MgCl_2_ for BT1636p and N4-ORF16p, respectively. Slides were rinsed to remove the enzyme and salts as described above for the serum sample. Samples were then dried in a desiccator.

#### Sample preparation for MALDI-MSI

Glycerol-free concentrated PNGase F (formulated internally) was applied to samples using an HTX M3+ (HTX Technologies LLC). 0.1 mg/mL PNGase F was applied in 15 passes with a nozzle temperature of 45°C, a nozzle height of 40 mm, a nozzle velocity of 1200 mm/min, a flow rate of 25 µL/min, a track spacing of 3 mm, and a nitrogen pressure of 10 psi. Following application, samples were incubated at 37°C in a humidity chamber for 2 hours and then dried in a desiccator for 10 minutes.

Samples were coated with α-cyano-4-hydroxycinnamic acid (CHCA) (Bruker) using an HTX M3+ sprayer (HTX Technologies LLC). A solution of 10 mg/mL CHCA in 70% acetonitrile, 0.1% trifluoroacetic acid, 20mM ammonium phosphate was applied in 3 passes with a nozzle temperature of 75°C, a nozzle height of 40 mm, a nozzle velocity of 1200 mm/min, a flow rate of 120 µL/min, a track spacing of 3 mm, and a nitrogen pressure of 10 psi.

### MALDI-MSI data acquisition and analysis

Images were acquired on a Bruker timsTOF flex mass spectrometer (Bruker Daltonics) in positive mode at a spatial resolution of 50 μm over the *m/z* range 700-4000 with a Funnel 1 RF of 500 Vpp, a Funnel 2 RF of 500 Vpp, a Multiple RF of 500 Vpp, a Collision Energy of 15 eV, a Collision RF of 4000 Vpp, a Transfer Time of 110 μs, a PrePulse Storage Time of 24 μs, and a laser power of 35%. A total of 600 laser shots were summed per pixel at 10kHz. Mass calibration was performed using Tune Mix (Agilent) and red phosphorus. Ion mobility was turned off for these experiments.

Data were analyzed using the SCiLS Lab software (Bruker, version 2025b 13.01.17192), and spectra were normalized to the total ion count. *N*-glycan peaks were manually identified through comparison to previously characterized *N*-glycans^6,10,21,22^.

### Functional metagenomic screening for sulfate-targeting enzymes

4-Methylumbelliferyl Galactose-3-SO_4_ (Glycouniverse, GU-CSYN-046) was used in a coupled fluorescence assay to screen for novel galactose-3-sulfatases as previously described^3^. Briefly, clones from a human gut microbiome metagenomic fosmid library were grown statically overnight at 37°C in 384-well plates with 50 µL Lennox Broth containing 12.5 μg/mL chloramphenicol and 1X autoinducing solution (Lucigen). To perform the screen, 50 µL of Y-PER™ lysis buffer (Thermo Scientific, 78990) supplemented with 120 μM of 4MU-Gal-3-SO_4_ and 1 U/mL of β1-3 Galactosidase (NEB, P0726) were added to each well. The plates were then incubated statically at 37°C for 48 hours. Fluorescence was read at λ_ex_=350 nm and λ_em_=445 nm after 1, 6, 24, and 48 hours using a SpectraMax Plus Microplate Reader (Molecular Devices). Clones were identified as hits if they exhibited fluorescence at least three standard deviations above background fluorescence of the mean fluorescence (excluding outliers). Fosmids from hit clones were sequenced on a RSII sequencer (Pacific Bioscience, USA). ORFs were predicted with MetaGeneMark^23^ and annotation of the predicted ORFs was performed using an accelerated profile HMM search^24^.

### Novel sulfatase characterization

#### Sulfatase activity on monosaccharides

To identify the metal ion cofactor requirement for N4-ORF16p and E8-ORF27p, 1 µM enzyme was incubated with 30 µM 4MU-Gal-3-SO_4_, 1 U/mL of β1-3 Galactosidase (NEB, P0726), 5 mM of MgCl_2_, CaCl_2_, CuSO_4_, NiSO_4_, MnSO_4_, FeSO_4_, CoCl_2_, or ZnSO_4_ in 50 mM Tris, pH 7.5. Reactions were performed in triplicate and incubated for 1h at 37°C. Fluorescence was read at λ_ex_=350 nm and λ_em_=445 nm using a SpectraMax Plus Microplate Reader (Molecular Devices).

To determine the optimal temperature for N4-ORF16p and E8-ORF27p, 1 µM enzyme and 0.1 mM galactose-3-SO_4_ (Toronto Research Chemicals, G155295) were incubated at temperatures ranging from 16 to 70°C. The optimal pH was determined by incubating 1 µM enzyme and 0.1 mM galactose-3-SO_4_, at the respective optimal temperatures in buffers ranging from pH 4.5 to 10.7 (from pH 4.5–5.5, 50 mM sodium acetate; from pH 6.0–7.0, 20 mM sodium phosphate; from pH 7.5–9, 50 mM Tris–HCl; pH 9.7, 20 mM N-cyclohexyl-3-aminopropanesulfonic acid; and pH 10.7, 50 mM carbonate buffer) in 10 μL final reaction volume. Reactions were performed in triplicate. After 1h incubation, samples were dried in a centrifuge concentrator (Vacufuge plus, Eppendorf) for 30 minutes. Dried reactions were then labeled with procainamide. The procainamide solution was prepared by dissolving 12 mg of procainamide (Abcam, ab120955) in 110 µL of 70% dimethyl sulfoxide/30% acetic acid. This solution and 25 µL water were then transferred to a tube containing 6 mg sodium cyanoborohydride (Sigma, 156159). 20 µL procainamide solution was added to each sample. Reactions were incubated for 1h at 65°C. Samples were diluted 1:10 in 100% acetonitrile and separated by UPLC using an ACQUITY UPLC glycan BEH amide column 130Å (2.1 × 150 mm, 1.7 μm) on a H-Class ACQUITY instrument (Waters). The injection volume was 3 μl. Solvent A was 50 mM ammonium formate buffer, pH 4.4, and solvent B was 100% acetonitrile. The flow rate was 0.4 mL/min and the gradient used was 0 to 35 min, 12 to 47% solvent A; 35 to 35.5 min, 47 to 70% solvent A; 35.5 to 36.0 min, 70% solvent A; 36 to 36.5 min, 70 to 12% solvent A; 36.5 to 40 min, 12% solvent A. Samples were kept at 4°C prior to injection and the column temperature was 40°C. Procainamide fluorescence was read at λ_ex_=308nm and λ_em_=359nm, with a data collection rate of 20 Hz.

Specificity of N4-ORF16p and E8-ORF27p was tested on the substrates listed in **Table S2** using the optimal reaction conditions identified above (**Table S3**). Following incubation with the sulfatases, samples were labeled with procainamide and analyzed by UPLC as described above.

#### Sulfatase activity on thyroglobulin N-glycans

*N*-glycans were released from human thyroglobulin (Sigma, T6830) with PNGase F (NEB, P0704). An aliquot containing 100 µg of thyroglobulin was incubated with 1X glycoprotein denaturing buffer and water to make a 100 µL total reaction volume. The glycoprotein was denatured at 100°C for 10 minutes. *N*-glycans were cleaved by adding 7 U/mL PNGase F, 1X glycobuffer 2, 1% NP-40 and water for a total reaction volume of 200 µL. The reaction was incubated at 37°C for 2 h and then cleaned up using a HyperSep Hypercarb SPE column (ThermoFisher 60106-302). Released *N*-glycans were labeled with 2-aminobenzamide (2-AB; Sigma, 76884) and excess 2-AB was removed through acetone precipitation. To confirm localization of the sulfate to galactose on these *N*-glycans, a sample was incubated with β1-3,4-galactosidase (NEB, P0746) which cannot cleave sulfated galactose. Samples were then analyzed by LC-MS. High-performance liquid chromatography was carried out on a ThermoFisher Vanquish system using fluorescence detection and separated using an ACQUITY UPLC Glycan BEH Amide (130 Å, 1.7 µm, 2.1 mm x 150 mm) column. The mobile phases used were A: 0.1% difluoro acetic acid in water and B: 0.1% difluoro acetic acid in acetonitrile. The column temperature was 40°C, flow rate 0.3 mL/min, and the gradient was: 70% B to 50% B from 0 to 35 min, 10% B from 35.5 to 38.5 min, with re-equilibration at 70% B from 39-45 min. 2-AB fluorescence was detected at λ_ex_=320 nm and λ_em_=420 nm. LC-MS/MS was performed using a ThermoFisher Orbitrap Eclipse in positive mode. Sulfated *N*-glycans were isolated through fractionation, incubated with N4-ORF16p or E8-ORF27p, and activity was analyzed by LC-MS as described above.

## RESULTS

### Integration of PGM-targeting enzymes into the MALDI-MSI analytical workffow

The primary aim of this study was to use new PGM-targeting enzymes in MALDI-MSI to structurally characterize *N*-glycan PGMs *in situ* and visualize their spatial distribution in biological samples. To accomplish this, three types of PGM-targeting enzymes were incorporated into a common MALDI-MSI *N*-glycan sample preparation and analysis workflow (**Fig 1**). In this type of analysis, tissues are sectioned, washed, and incubated with the enzyme PNGase F to release *N*-glycans after which they are imaged by MALDI-TOF^10^. In the present study, adjacent tissue sections were first treated with enzymes acting on a specific PGM prior to *N*-glycan release by PNGase F. When the intensity of an MS peak decreases following treatment with a PGM-targeting enzyme of known specificity, the sugar residue, the position of the PGM modification on the sugar, and the spatial position of the modified *N*-glycan in the tissue can be determined. In principle, different enzyme specificities could be used to discriminate different types of *N*-glycan modifications, providing a holistic view of the composition and spatial distribution of PGMs in the *in situ N*-glycome. In this study, we establish this method and verify the suitability of three PGM-targeting enzyme classes, a phosphodiesterase, a glycosyl hydrolase, and sulfatases, for this type of analysis, using invertebrate tissue containing phosphorylcholine-modified *N*-glycans, and both mammalian serum and mammalian tissues containing different types of sulfated *N*-glycans.

**Figure 1.**
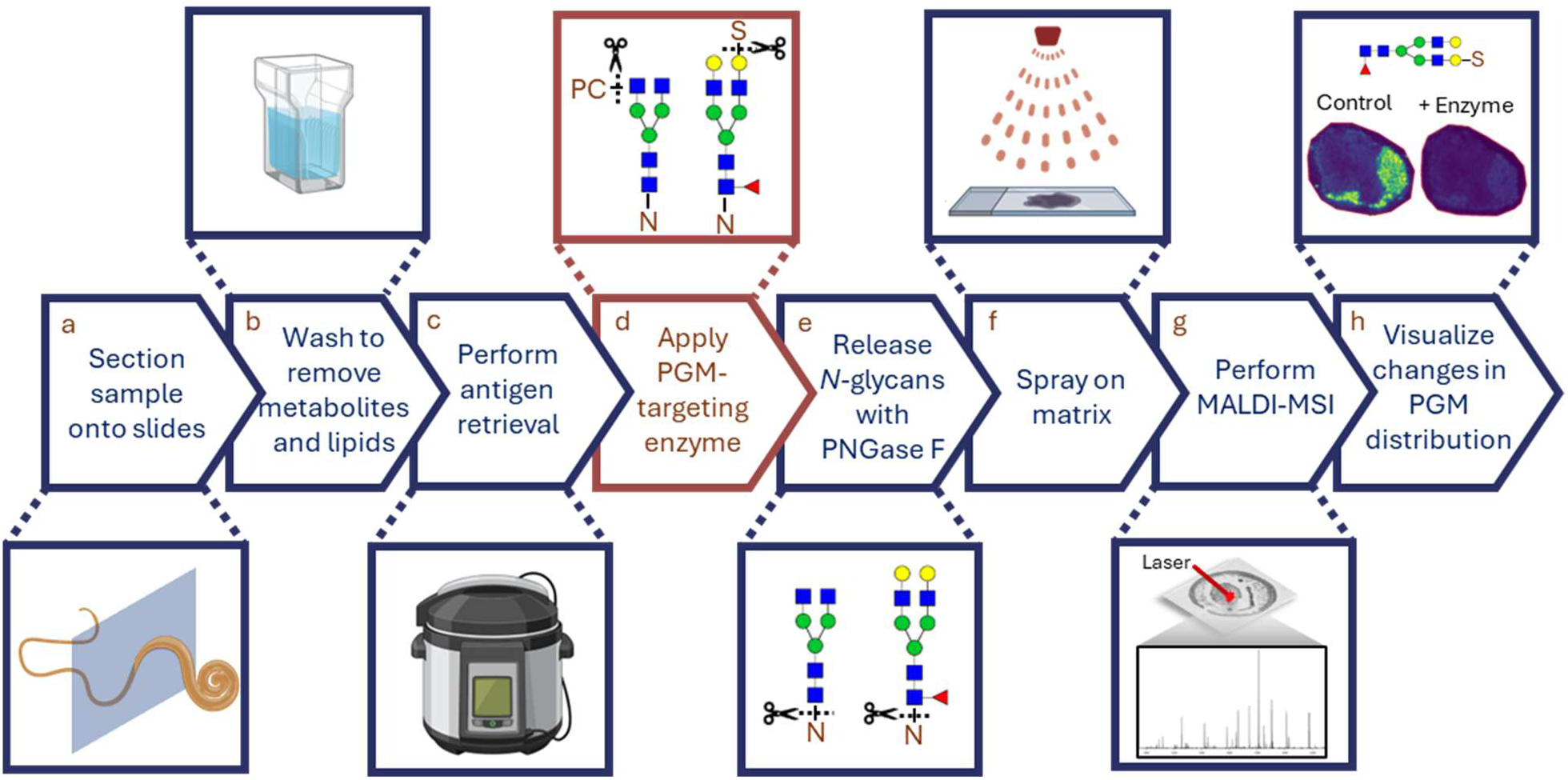
Incorporation of post-glycosylation modifying enzymes into the *N*-glycan MALDI-MSI workflow. **a)** Samples are first embedded or frozen, sectioned, and mounted on MALDI compatible slides. **b)** Metabolites and lipids are then removed through a series of washes **c)** Antigen retrieval is then performed. **d)** Samples are then treated with a PGM-targeting enzyme specific to the modification of interest. **e)** PNGase F is then sprayed onto the sample to liberate *N*-glycans from glycoproteins. **f)** MALDI matrix is then sprayed onto the sample and **g)** MALDI-MSI is performed. **h)** Comparison of samples treated with or without a PGM targeting enzyme allows for visualization of changes to PGM distribution. Figure made in part with Biorender.com.

### GlcNAc-C phosphodiesterase enables visualization of phosphorylcholine in roundworm N-glycans

The roundworm *Ascaris suum* is an intestinal parasitic nematode that produces *N*-glycans modified by the zwitterion phosphorylcholine (PC)^22^. PC-modified glycans play an important immunomodulatory role during infections for many parasitic nematodes^25–30^. To confirm the presence of *N*-glycans with GlcNAc-6-PC and visualize their *in situ* localization in *A. suum* sections, a GlcNAc-6 phosphodiesterase (termed GlcNAc-6-PDase) was used in the MALDI-MSI workflow described above^14^. This enzyme was previously shown to have remarkable selectivity for GlcNAc-6-PC and no activity on mannose-PC, another common PGM in some invertebrate *N*-glycans, including those from nematodes^14^. When *A. suum N-*glycans were released with PNGase F, several *N-*glycans that contain PC were seen localized either to the cuticle, muscle cell bags, or ovaries, but not in the *Ascaris* intestine or uterus (**Fig 2**). These data both confirm the presence of GlcNAc-6-PC in these *N*-glycans and show their spatial distribution in worm tissue. It also demonstrates that PGMs can be enzymatically manipulated in a highly complex *in situ* environment for MALDI-MSI.

**Figure 2.**
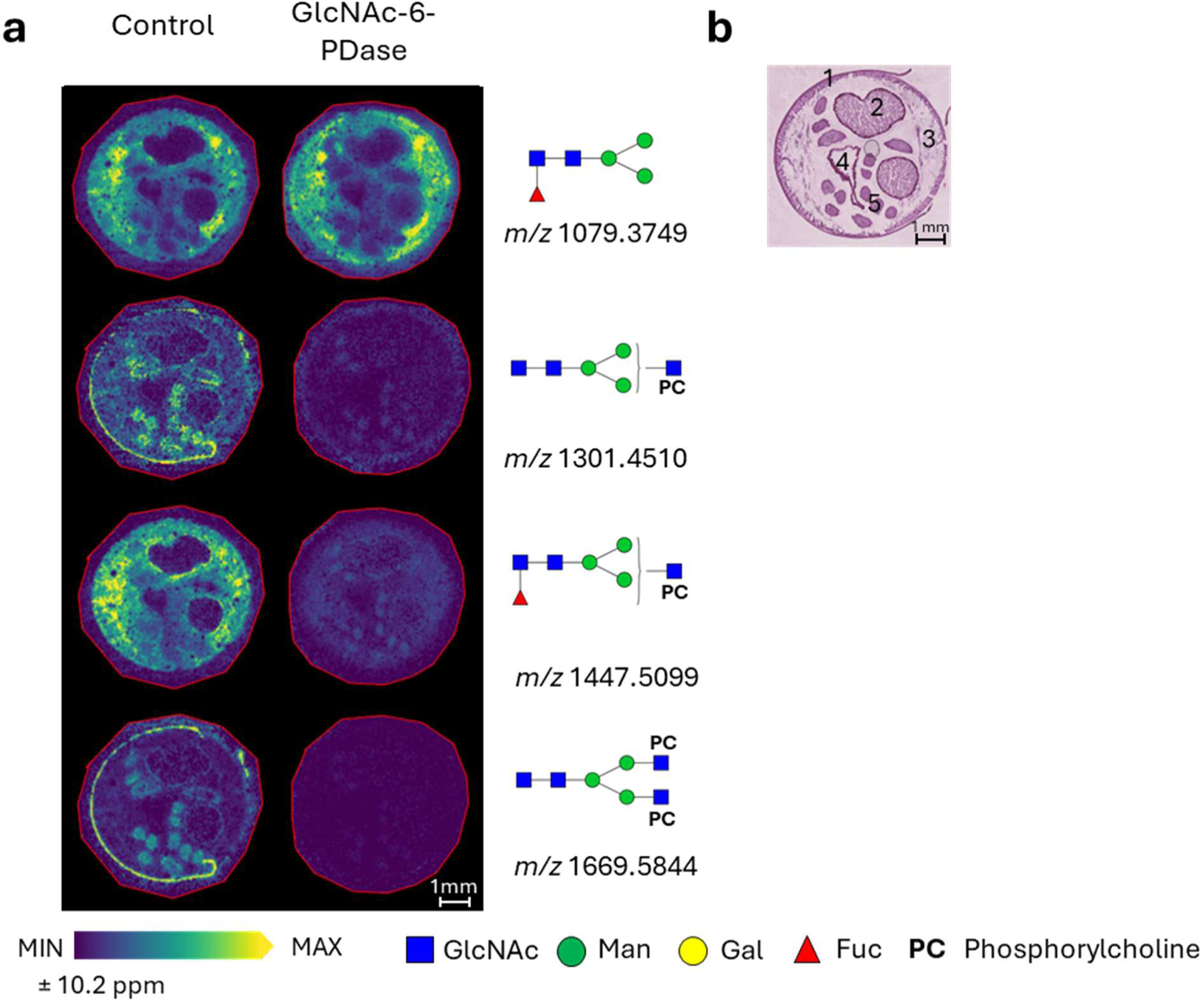
Use of GlcNAc-6-PDase to visualize distribution of phosphorylcholine in *A. suum*. **a)** MALDI imaging of neutral and zwitterionic *N-*glycans (Hex3dHex1HexNAc2 + 1Na *m/z* 1079.3749; Hex3HexNAc3 + 1PC + 1Na *m/z* 1301.4510; Hex3dHex1HexNAc3 + 1PC + 1Na *m/z* 1447.5099; Hex3HexNAc4 + 2PC + 1Na *m/z* 1669.5844 within a ± 10.2 ppm mass error) present in *A. suum*. Serial sections were treated with or without GlcNAc-6-PDase followed by treatment with PNGaseF to release *N-*glycans. Key: *N-*acetyl glucosamine (GlcNAc), mannose (Man), galactose (Gal), fucose (Fuc), phosphorylcholine (PC). **b)** HCE stain of a serial section. Key: 1 – cuticle, 2 – uterus with eggs, 3 – muscle, 4 – intestine, 5 – ovary.

### Use of sulfate-targeting enzymes to characterize N-glycans in human serum

Sulfation is a common PGM found in nearly all organisms on a wide diversity of glycans, including glycosphingolipids, *N*- and *O*-linked glycans, glycosaminoglycans, and marine exopolysaccharides^3,31^. As such, glycan sulfation influences a wide variety of processes, including altering the structural stability of surface adhesion, determining the host-specificity of symbiotic relationships, and modulating host-pathogen interactions^32–35^. Accordingly, changes in sulfation patterns have significant effects on glycan function. For instance, decreases in dermatan sulfate sulfation impede collagen biosynthesis causing the congenital connective tissue disease, Ehlers-Danlos syndrome^36^. As with other PGMs, glycan sulfation continues to be understudied despite its ubiquitous importance due to its structural complexity. For example, in structural studies of protein *N*- and *O*-glycosylation, sulfation has been observed on the 3^rd^, 4^th^, or 6^th^ carbon of mannose, galactose, GlcNAc and *N*-acetylgalactosamine (GalNAc)^3^. Thus, to understand the distribution of sulfation *in situ*, enzymes that both discriminate between types of sulfated sugar and identify the linkage of the sulfate modification will be imperative. Ultimately, these enzymes must also function efficiently in a highly complex extracellular environment to be suitable for MALDI-MSI.

We established workflow conditions using a sulfate-targeting glycoside hydrolase with bulk serum glycoproteins. Serum glycoproteins were examined because the sample would be less complex than the surface of a sectioned tissue, and *N*-glycan sulfation has been described for a subset of serum glycoproteins^6,37^. For example, numerous *N*-glycans released from serum immunoglobulins are modified with GlcNAc-6-SO_4_^3,6^ In this experiment, we utilized the enzyme *N*-acetyl-glucosamine-6-SO_4_-hexosaminidase (F10-ORF19p) to remove GlcNAc-6-SO_4_ from serum glycoprotein *N*-glycans enzymatically ^3^. Non-reactive human serum was spotted onto amine-reactive slides that covalently bind proteins by their primary amine groups. Because this hexosaminidase only acts on terminal GlcNAc-6-SO_4_ residues, the sample was first treated with Neuraminidase A and β-galactosidase to sequentially remove sialic acid and galactose residues and expose terminal GlcNAc-6-SO_4_. The samples were then incubated with GlcNAc-6-SO_4_-hexosaminidase, after which, *N*-glycans were released with PNGase F, and the samples were analyzed by MALDI MS (**Fig 3, Fig S1**). There was a marked decrease in signal for several sulfated *N*-glycans and a corresponding increase of *N*-glycans where GlcNAc-6-hexosaminidase had removed terminal GlcNAc-SO_4_, consistent with the known substrate requirements of F10-ORF19p. These data confirm the presence of GlcNAc-6-SO_4_ on human serum *N*-glycans and demonstrate the utility of PGM-targeting enzymes to address a second PGM, sulfation.

**Figure 3.**
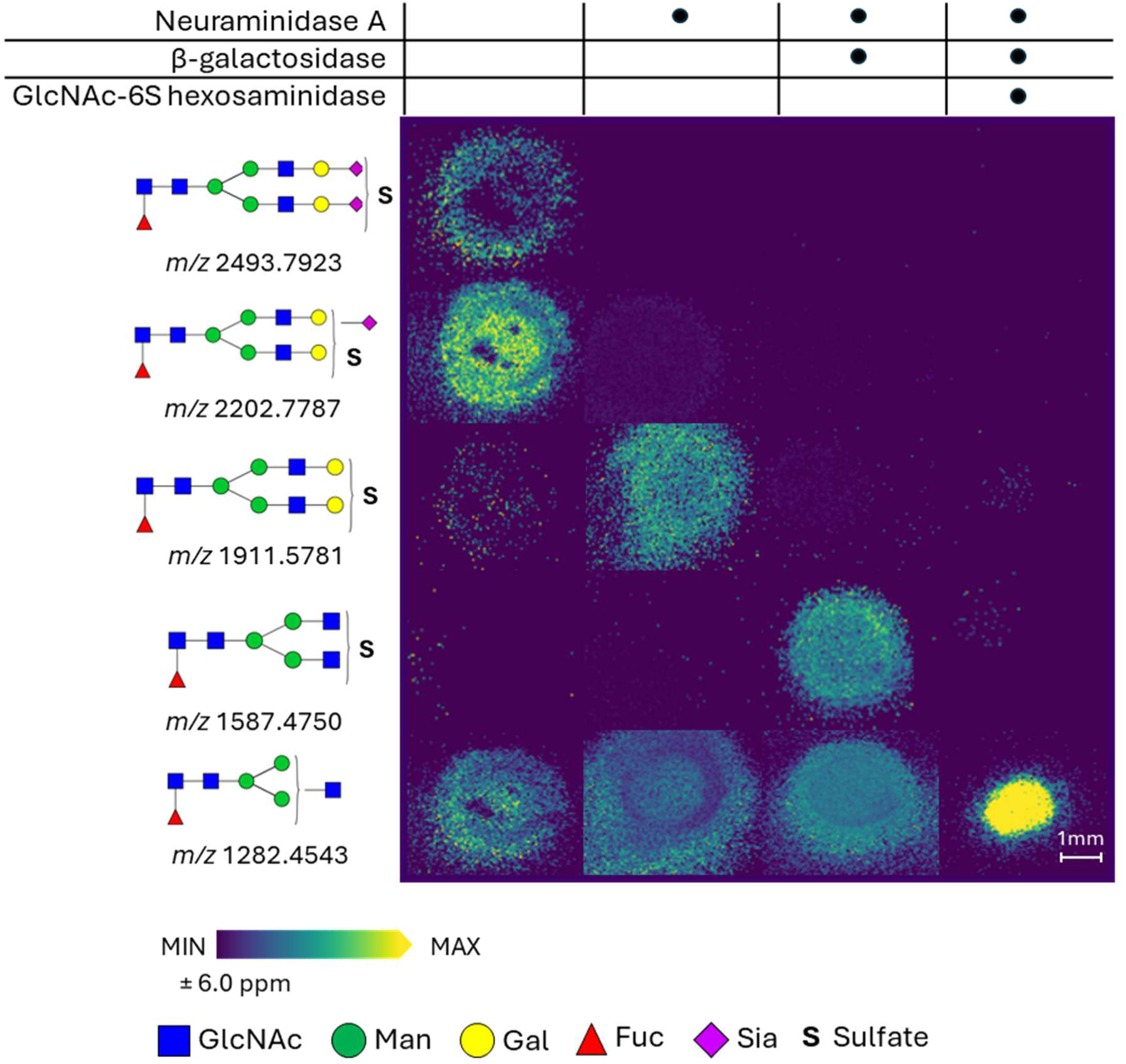
Use of GlcNAc-6-sulfate hexosaminidase to visualize glycan sulfation in human serum. MALDI imaging of exoglycosidase treated *N-*glycans. (Hex5dHex1HexNAc4NeuAc2 + 1SO_4_ + 2Na *m/z* 2493.7923; Hex5dHex1HexNAc4NeuAc1 + 1SO_4_ + 2Na *m/z* 2202.7787; Hex5dHex1HexNAc4 + 1SO_4_ + 2Na *m/z* 1911.5781; Hex3dHex1HexNAc4 + 1SO_4_ + 2Na *m/z* 1587.475; Hex3dHex1HexNAc3 *m/z* 1282.4543 within a ± 6.0ppm mass error). Serum was spotted on amine reactive slides and separate spots of serum were treated with Neuraminidase A and β-galactosidase to expose GlcNAc residues followed by treatment with a GlcNAc-6-SO_4_ specific hexosaminidase (F10-ORF19p, Chuzel et al., 2021). *N-*glycans were then released using PNGase F. Key: *N-*acetyl glucosamine (GlcNAc), mannose (Man), galactose (Gal), fucose (Fuc), sialic acid (Sia), sulfate (S).

### Discovery and characterization of new sulfate-targeting enzymes

A more comprehensive arsenal of sulfate-acting enzymes will be needed to systematically decode the sulfated *N*-glycome *in situ*. Although the number of characterized sulfate-targeting enzymes has increased in recent years (**Table 1**), there are still sulfate positions that cannot be targeted by known enzymes. Additionally, enzymes with broader substrate recognition will enable a “funnel” approach for characterization. In this approach a few broad specificity enzymes would initially be applied to the sample, and once the nature of the modification is identified more specific enzymes could be applied to fully elucidate glycan structure. This approach would facilitate higher-throughput analysis of sulfated glycans and minimize the amount of sample needed for characterization. However, the discovery of additional sulfatase specificities is needed.

**Table 1.**
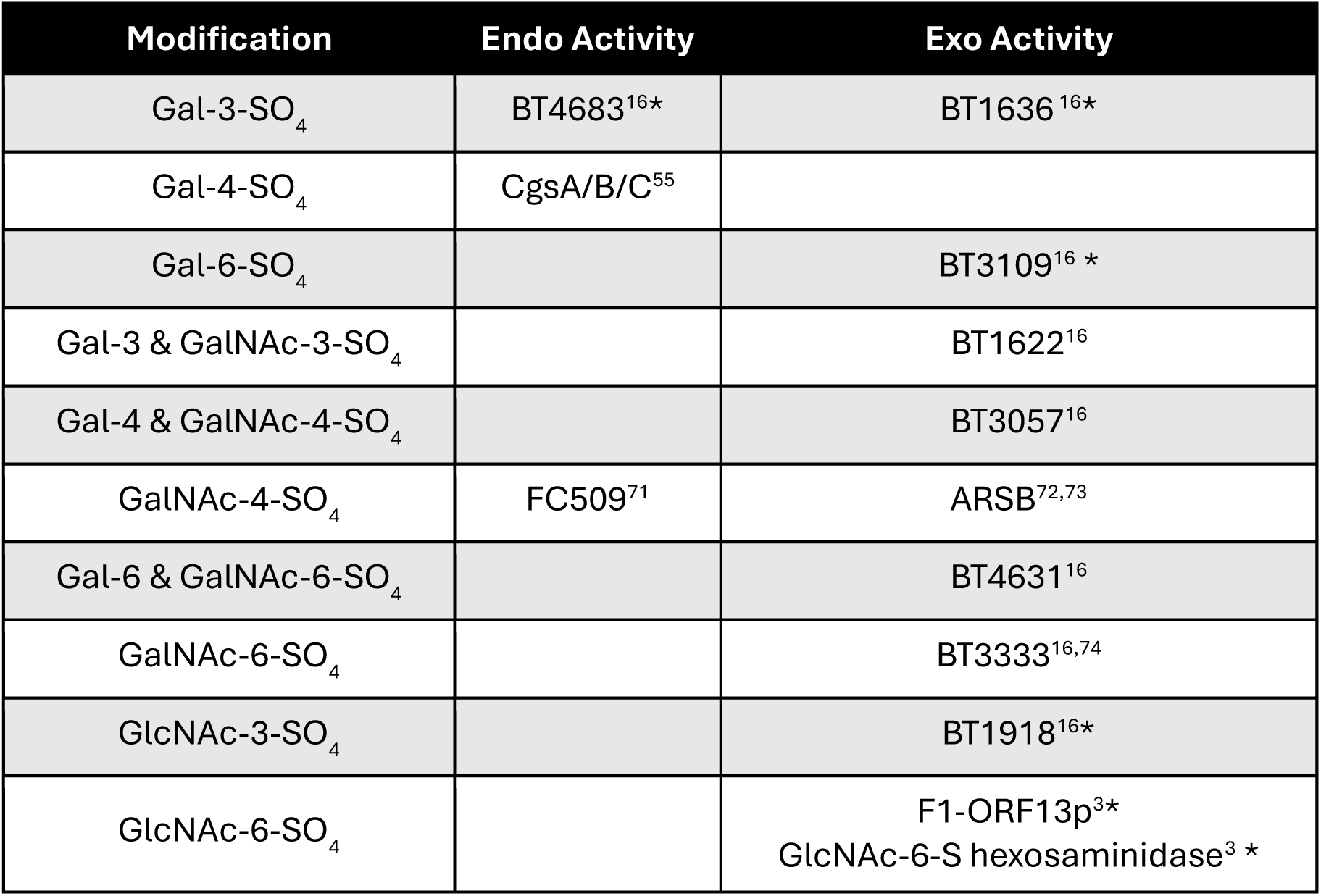
Sulfate targeting Toolbox. Summary of previously characterized enzymes that can be used to study sulfated glycans. Examples of each type of activity have been given, additional enzymes have been characterized for several of these modifications. *Activity independently verified in-house.

To identify new sulfate-targeting enzymes, a functional metagenomic screen was performed for enzymes that act on galactose-3-SO_4_. We reasoned that screening for enzymes that act upon sulfated galactose may identify others with broader specificities. The assay used here was adapted from our previous screen that identified several sulfate-targeting enzymes specific to GlcNAc-6-SO_4_^3^. Briefly, 4-methylumbelliferone (4-MU)-labeled galactose-3-SO_4_ was incubated with cell lysates from a human gut microbiome fosmid library. When a sulfatase expressed from the fosmid removes the sulfate from the substrate an exogenous β-galactosidase provided in the assay mixture hydrolyzes 4MU-β-Gal and generates fluorescence (**Fig 4A**). A clone was considered a “hit” if it produced a signal greater than 3 standard deviations above the mean background fluorescence. A total of 5,376 clones were screened and produced 9 hits. Upon rescreening with the same assay, 5 of the clones exhibited a reproducible signal, exhibiting signal greater than 6 stand deviations above background (**Fig S2A**). Lysates from these 5 clones were additionally assayed with different substrates and assay conditions: i) 4MU-Gal-3-SO_4_ (without exogenous β-galactosidase), to identify β-galactosidases that are not inhibited by galactose-3-sulfation, ii) 4MU-Gal to detect β-galactosidase activity, and iii) 4MU-SO_4_ to detect indiscriminate sulfatase activity (**Fig S2B**). Three of the clones exhibited β-galactosidase activity, an activity supported by the presence of sulfatase and β-galactosidase genes in those sequences. Annotation of the DNA sequence from each of the 5 candidate fosmid clones revealed overlap in their insert sequences, resulting in identification of 2 unique sulfatase candidates (**Fig S2C**). The two candidates, herein termed N4-ORF16p and E8-ORF27p, belong to two different sulfatase subfamilies. N4-ORF16p is the second sulfatase characterized from subfamily S1_46, with the previously described sulfatase acting on GlcNAc-3-SO_4_^16,31,38^. E8-ORF27p is a member of subfamily S1_65 and is the first member of this subfamily to be functionally characterized. Sulfatase activity of heterologously expressed and purified candidates was confirmed by incubating each recombinant protein with 4MU-Gal-3-SO_4_ in the presence of exogenous β-galactosidase (**Fig S3**)^31^.

**Figure 4.**
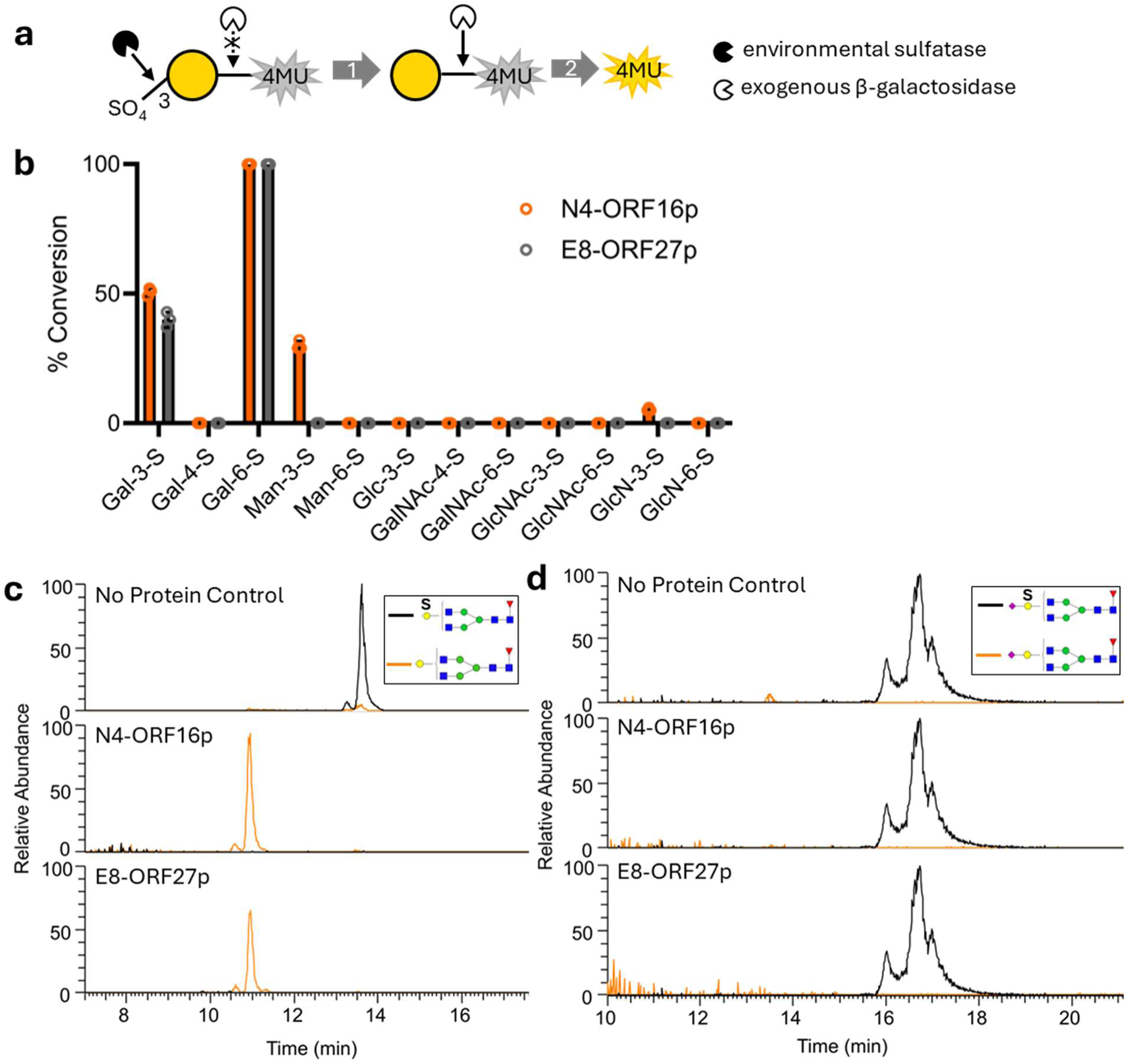
Functional metagenomic screening for sulfatases that act on galactose-3-SO_4_. **a)** Coupled assay used for sulfatase screening. The substrate used was galactose-3-sulfate linked to the fluorophore 4-methylumbelliferone (4MU). When incubated with a metagenomic library clone encoding a sulfatase that can remove the sulfate, the exogenous galactosidase cleaves 4MU, generating fluorescence. **b)** Sulfatase specificity was determined using procainamide labeled sulfated monosaccharides and percent conversion to de-sulfated form was analyzed by UPLC. Data presented as mean and standard deviation, N = 3. **c)** Sulfatase activity was evaluated on *N-* glycans isolated from human thyroglobulin, labeled with 2-AB, and treated with β-galactosidase (**Fig S5**). Isolated *N-*glycans tested contained either a terminal sulfated galactose (Hex4dHex1HexNAc4 + 1SO_4_ *m/z* 913.322; Hex4dHex1HexNAc4 *m/z* 873.3428 ± 10.0ppm mass error) or **d)** internal sulfated galactose (Hex4dHex1HexNAc4NeuAc1 + 1SO_4_ *m/z* 1058.864; Hex4dHex1HexNAc4NeuAc1 *m/z* 1018.89 ± 10.0ppm mass error). Extracted ion chromatograms are normalized to the sulfated *N-*glycan signal in the no enzyme controls.

Since sulfatases frequently exhibit activity on multiple substrates, the substrate specificities of N4-ORF16p and E8-ORF27p were evaluated using a panel of sulfated monosaccharides (**Table S2**). Reactions were carried out using the optimal pH, temperature, and metal cofactors empirically determined for each sulfatase (**Fig S4**). Activity was quantified as the percent conversion of a sulfated monosaccharide to its de-sulfated form after one hour of incubation (**Fig 4B**). The two sulfatases exhibited overlapping, but distinct specificities. Both enzymes act on galactose-3-SO_4_ and galactose-6-SO_4_, with an apparent preference for the latter, while N4-ORF16p also acted on mannose-3-SO_4_, and to a far lesser extent glucosamine-3-SO_4_. These broader specificities differentiate these sulfatases from the previously described galactose-3-sulfatases which act solely on galactose-3-SO_4_, either exclusively on internal residues (BT4683p) or exclusively on terminal residues (BT1636p)^16^, expanding the types of enzymatic tools at our disposal.

N4-ORF16p and E8-ORF27p were evaluated for their ability to act on more complex sulfated *N*-glycan substrates. The human glycoprotein thyroglobulin was previously shown to contain *N*-glycans with sulfated galactose residues ^39^. *N*-glycans were released from thyroglobulin, treated with β-galactosidase, and internally or terminally sulfated *N*-glycans were isolated via HPLC fractionation (**Fig S5**). The sulfated *N*-glycans were then incubated with N4-ORF16p or E8-ORF27p and analyzed by LC-MS. Both enzymes removed sulfate from the *N*-glycan with a sulfated terminal galactose but not from a glycan with a sulfated internal galactose (**Fig 4CD**).

### Application of novel sulfatases in MALDI-MSI of cancerous human thyroid tissue

Enzymes that site-specifically remove sulfate from sugars were used in MALDI-MSI to characterize poorly understood sulfated *N*-glycans. Some human thyroid glycoproteins contain sulfated galactose, however, the position of the sulfate modifications is unknown^39^. In this experiment, parallel sulfatase treatment was performed on serial sections of cancerous human thyroid tissue ahead of spatial imaging using MALDI-MSI. The sulfatases used have different specificities to discriminate between sulfated positions *in situ*. First, the newly discovered broad specificity sulfatase E8-ORF27p (exo-galactose-3/6-sulfatase) was applied to determine the nature of galactose sulfation in this sample (**Fig 5A, Fig S6**). A clear decrease in signal for sulfated *N*-glycan structures was observed, indicating that galactose is modified either on the 3^rd^ or 6^th^ carbon. A serial section of tissue was then treated with an exo-galactose-3-sulfatase (BT1636p)^16^ to further refine the position of the sulfate. The same decrease in signal was observed, indicating that all the sulfated *N*-glycans imaged contain galactose-3-SO_4_. Additional serial sections were treated with an exo-galactose-6-sulfatase (BT3109p)^16^ and an endo-galactose-3-sulfatase (BT4683p)^16^. These two enzymes had no effect on the sulfated *N*-glycans, corroborating the observation that terminal galactose-3-SO_4_ predominates in this tissue. This observation is further supported by a recent study that observed galactose-3-SO_4_ sulfation on *N*-glycans extracted from cancerous human thyroid tissue^21^.

**Figure 5.**
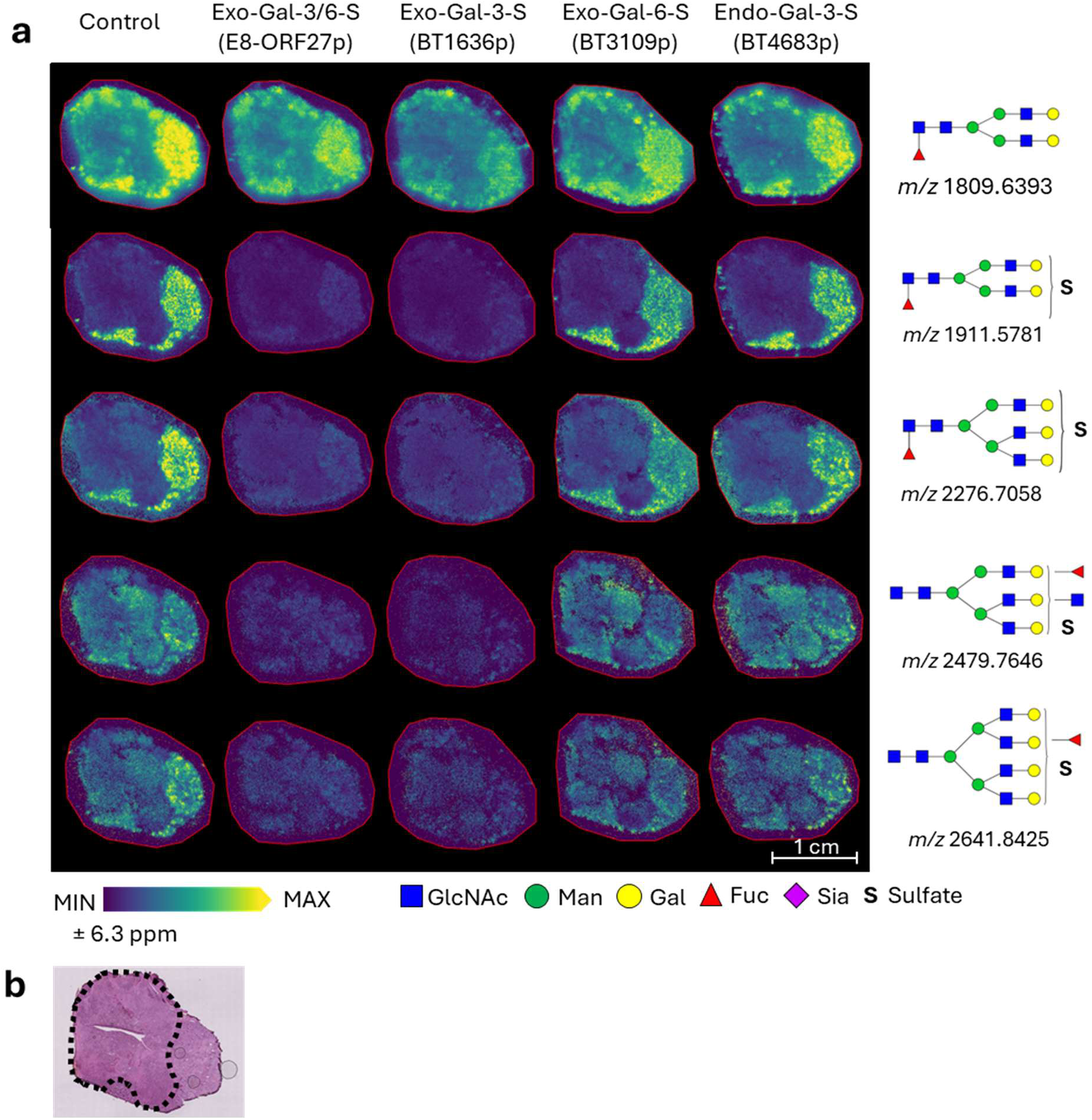
Use of sulfatases to visualize the identity and distribution of sulfated *N*-glycans in cancerous human thyroid tissue. **A)** MALDI imaging of sulfated *N*-glycans (Hex5dHex1HexNAc4 + 1Na *m/z* 1809.6393; Hex5dHex1HexNAc4 + 1SO_4_ + 2Na *m/z* 1911.5781; Hex6dHex1HexNAc5 + 1SO_4_ + 2Na *m/z* 2276.7058; Hex6dHex1HexNAc6 + 1SO_4_ + 2Na *m/z* 2479.7646; Hex7dHex1HexNAc6 + 1SO_4_ + 2Na *m/z* 2641.8425 within a ± 6.3 ppm mass error) present in cancerous human thyroid tissue. Serial sections were treated with either an endo-galactose-3/6-sulfatase (E8-ORF27p), an exo-galactose-3-sulfatase (BT1636p), an exo-galactose-6-sulfatase (BT3109p) or an endo-galactose-3-sulfatase (BT4683p) (Luis et al., 2021) followed by treatment with PNGaseF to release *N*-glycans. **B)** HCE stain of a neighboring section, cancerous region is outlined in black. Key: *N*-acetyl glucosamine (GlcNAc), mannose (Man), galactose (Gal), fucose (Fuc), sialic acid (Sia), sulfate (S).

These data demonstrate that both broadly acting and highly specific PGM-targeting enzymes can be used together in MALDI-MSI tissue imaging to determine the nature and spatial distribution of PGM chemical marks in the *N*-glycome. Furthermore, six of the sulfated *N*-glycans imaged in this experiment, were observed only in the healthy region of the tissue. While this was a singular compelling experiment, it highlights the potential importance and impact of determining spatial PGM information in researching clinical samples (**Fig 5B**).

## DISCUSSION

In this study, we established the use of PGM-targeting enzymes in MALDI-MSI for rapid characterization and mapping of modified *N*-glycans *in situ*. This approach simultaneously elucidates the PGM-containing sugar, the position of the modification, and the spatial distribution of the modified *N*-glycan on the surface of a tissue. We first illustrated the approach using a highly specific GlcNAc-6-phosphodiesterase to localize PC-containing *N*-glycans in the invertebrate *A. suum*. Next, we showed localization of sulfated mammalian *N*-glycans harboring GlcNAc-6-SO_4_. This experiment utilized a sulfated sugar-specific glycoside hydrolase (GlcNAc-6-SO_4_ hexosaminidase) to visualize *N*-glycans having GlcNAc-6-SO_4_ in their outer arm. Finally, following the discovery of sulfatases with novel broad specificities, a series of sulfatases were used to identify and visualize the localization of sulfated *N*-glycans in cancerous human thyroid tissue. This study shows the first use of enzymes to achieve linkage-level resolution of PGMs in MALDI-MSI and enables exploration of spatially relevant PGM changes in the *N*-glycome that may correlate with disease.

The work presented in this study provides a methodological framework for visualizing the spatial distribution of PGMs in tissues. In the case of mammalian glycan sulfation, changes have been observed in association with several diseases. For instance, increases in *O-*glycan sulfation were observed in ovarian and breast cancers^40,41^. Changes in glycosaminoglycan (GAG) sulfation have been associated with numerous cancers as well as Alzheimer’s disease^42–45^. In each of these cases, sulfation states were analyzed through glycomics of homogenous samples. Additionally, recent work to develop a comprehensive understanding of tissue and disease-specific biomarkers have largely excluded PGMs since verification of the modifications would greatly decrease the throughput of these analyses^12,46–48^. This method enables analysis of changes to PGM patterns between tissues and disease states. Previous studies identifying changes of PGMs in disease were dependent on homogenous samples. Direct visualization of PGM spatial distribution, facilitated by studies such as ours, will help advance our understanding of cellular function and associated pathologies. Visualizing the distribution of sulfated glycans may provide additional insights into both the molecular nature of the disease and enable defining the borders of diseased tissue, as suggested by our analysis of cancerous thyroid tissue (**Fig 5, Fig S6**).

This study specifically addressed modification of *N*-glycans primarily because methods for their release and analysis by MALDI-MSI are well established^10^. Recent work has also demonstrated feasibility of imaging glycosphingolipids using MALDI-MSI, making them amenable to this type of analysis^49–51^. However, rapid analysis of other glycan classes, such as *O*-glycans, remains limited. Furthermore, glycosaminoglycans and exopolysaccharides are composed of repeating saccharide units and frequently contain multiple PGMs per repeat, significantly complicating their analysis^52–54^. As new methods are developed and the classes of glycans accessible through MSI are expanded, PGM-targeting enzymes can be implemented into any sample preparation workflow. Additionally, PGM-targeting enzymes can still be utilized as part of traditional exoglycosidase digestion analyses of modified glycans in homogenous samples.

To extend the application of PGM visualization with MSI, the toolbox of well characterized PGM targeting enzymes needs to be expanded. Recent studies elucidating novel sulfate-targeting enzymes with distinct specificities have established a solid foundation for the enzymatic toolbox needed for these experiments^3,16,31,55–65^. However, *N-*glycan sulfation is complex, and the field is still limited by the types of sulfation that can be enzymatically targeted. For example, there are sulfated glycans that are not recognized by any known enzymes and most described sulfate-targeting enzymes act only on terminal or internal residues (**Table 1**). Since there are no known endo-acting GlcNAc-6-SO_4_ targeting enzymes, it was necessary for our experiment to first expose the sulfated residues on serum *N*-glycans through a series of enzymatic digestions (**Fig 3**). While this approach was effective, it increases the time and reagents needed for sample preparation. In addition to sulfation, certain sugar-specific deacetylases and phosphatases could in principle also be used to study PGMs with MSI, however, there are limited specificities described for these enzyme classes^66,67^. Finally, while sulfatases and other carbohydrate-active enzymes are easily recognized through bioinformatics, it is not yet possible to predict the substrate specificity of these enzymes^67,68^. Thus, function-based enzyme screening using methods like functional metagenomics, a sequence agnostic method of screening for specific enzymatic activities, will continue to be a valuable enzyme discovery approach^3,14,69^.

There are additional opportunities for optimization to increase throughput and minimize the amount of sample needed for MSI analysis of PGMs. Running each PGM-targeting enzyme condition separately quickly multiplies the amount of instrument time needed, which is not sustainable for high-throughput applications. Fortunately, for homogenous samples that do not require localization data (*e.g.* serum samples), multiple enzymatic treatments can be queried on a single slide using a multi-well slide manifold. For samples requiring localization information, this method requires serial tissue sections for each condition tested, increasing the amount of sample needed for full characterization. This may not be feasible for precious clinical samples. To reduce the sample and time required, enzymes with broader specificities, such as N4-ORF16p and E8-ORF27p that act on both Gal-3-SO_4_ and Gal-6-SO_4_ can be deployed to narrow down structural possibilities, instead of querying each possible position individually (**Fig 5A, S6**). Faster characterization of glycan modifications will become possible as more PGM enzymes with broader specificities are described. Additionally, multiple section types could be manually mounted to the same slide for enzyme treatment. Alternatively, the use of tissue microarrays will enable screening for disease biomarkers in many samples in parallel further reducing instrument time required^70^.

Chemically modified glycans are abundant in the biosphere but remain cumbersome to characterize and study. By applying specific PGM-targeting enzymes to a MALDI-MSI workflow, we successfully enabled rapid determination of the presence of modified glycans, the sugar residue and position of specific modifications, and their localization in tissues. Incorporation of PGM-targeting enzymes into MSI studies will increase the accessibility of studying modified glycans and produce more structurally comprehensive glycomic analyses. Insight into when and where PGMs are present will enable a better understanding of biological functions as well as enable discovery of PGM-targeting biomarkers.

## AUTHOR CONTRIBUTIONS

J.E.D., C.H.T., and R.R.D. conceived of the study. J.E.D., L.C., E.E.E, G.G., H.B., A.H., S.L.F., and V.G. performed experimentation. J.E.D., L.C., E.E.E, and A.H. analyzed data. R.J.M. provided the *A. suum* samples. J.E.D. wrote the original draft.; J.E.D., L.C., E.E.E., A.H., S.L.F., R.R.D., J.M.F., and C.H.T. reviewed and edited the manuscript. C.H.T, J.M.F., and R.R.D. supervised the project. All authors read and approved the final manuscript.

## DATA AVAILABILITY

The datasets presented in this study can be found at https://glycopost.glycosmos.org/, GPST000589 or are available upon reasonable request to the authors.

## ACKNOWLEDGEMENTS

The authors thank Don Combs and New England Biolabs for basic research funding. This study is supported in part by the NIH, through the National Institute of Allergy and Infectious Diseases grants R01AI047194, R01AI155413 to R.J.M. and the E. A. Benbrook Foundation for Pathology and Parasitology.

## Notes

http://doi.org/10.50821/GLYCOPOST-GPST000589

### Competing Interest Statement

The authors have declared no competing interest.

