## Supplemental Figures for "Application of post glycosylation modifying enzymes for mass spectrometry imaging of modified *N*-glycans *in situ*"

| | |
|---|---|
| pJS119K_ORF16_Fwd | TAAGCTTAGGAGGTTAACATATGGATAAACATCCTAATATTATCC |
| pJS119K_ORF16_Rev | AAAACAGCCAAGCTGAATTCtcagtgatggtgatggtgatgTAACACCTCGGTGTTCTG |
| pJS119K_ORF27_Fwd | TAAGCTTAGGAGGTTAACATATGGCATATAATATCATTTTCTATTTCAGTGAC |
| pJS119K_ORF27_Rev | AAAACAGCCAAGCTGAATTCtcagtgatggtgatggtgatgGTCGGTGATAATGGGGCG |
| pJS119K_BT1636_Fwd | TAAGCTTAGGAGGTTAACATATGCAGAAAAACAACACAAAG |
| pJS119K_BT1636_Rev | AAAACAGCCAAGCTGAATTCtcagtgatggtgatggtgatgTTTCTTCTCCGGTAGTGTAAC |
| pJS119K_BT3109_Fwd | TAAGCTTAGGAGGTTAACATATGTCACAGCAGGTAGAAG |
| pJS119K_BT3019_Rev | AAAACAGCCAAGCTGAATTCTTAGTGATGGTGATGGTG |
| pJS119K_BT4683_Fwd | TAAGCTTAGGAGGTTAACATATGCAGGTAAGTACTCCTC |
| pJS119K_BT4683_Rev | AAAACAGCCAAGCTGAATTCtcagtgatggtgatggtgatgTTCTATTCCGCTTTGATC |
| BT3109_S57C_Fwd | TTCGGCTACT*tgc*ACCCCCAGTC |
| BT3109_S57C_Rev | GTAGCATATCCATTTGTGAAACAAAC |

**Supplemental Table 1.** Primers used in this study. Overlap with the cloning vector indicated by underlined text, tags inserted indicated by lowercase text, and bases changed in mutagenesis indicated by italicized text.

| | |
|---|---|
| Galactose-3-$SO_4$ | Toronto Research Chemicals (G155295) |
| Galactose-4-$SO_4$ | Dextra (G0012) |
| Galactose-6-$SO_4$ | Toronto Research Chemicals (G155298) |
| Mannose-3-$SO_4$ | GlycoUniverse (GU-CSYN-2375); custom synthesis) |
| Mannose-6-$SO_4$ | SantaCruz (sc-257290) |
| GalNAc-4-$SO_4$ | Dextra (G1054) |
| GalNAc-6-$SO_4$ | SantaCruz (sc-295652) |
| GlcNAc-3-$SO_4$ | GlycoUniverse (GU-CSYN-2378; custom synthesis) |
| GlcNAc-6-$SO_4$ | Biosynth (MA00759) |
| Glucose-3-$SO_4$ | Biosynth (FG146915) |
| Glucosamine-3-$SO_4$ | Sigma (11631) |
| Glucosamine-6-$SO_4$ | Dextra (G1002) |

**Supplemental Table 2.** Sulfated monosaccharides used in this study.

| | N4-ORF16p | E8-ORF27p |
|---|---|---|
| pH | Tris, pH 8.0 | Tris, pH 7.5 |
| Metal Cofactors | 5 mM $MgCl_2$ | 5 mM $MnSO_4$ |
| Temperature | 50°C | 22°C |

**Supplemental Table 3.** Optimal reaction conditions identified for novel sulfatases.

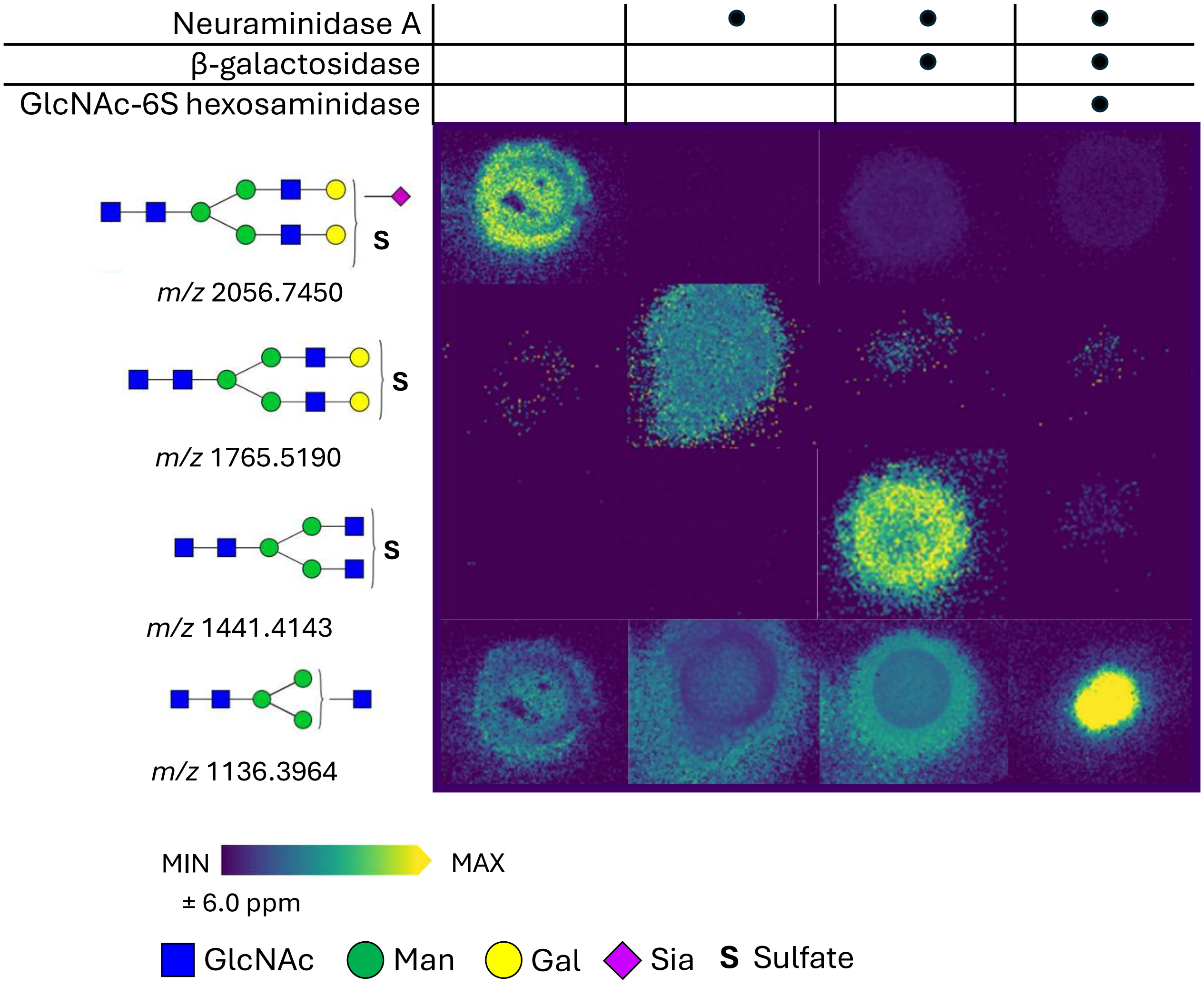


**Supplemental Figure 1. Additional example of the use of GlcNAc-6-sulfate hexosaminidase to visualize glycan sulfation in human serum**. MALDI imaging of exoglycosidase treated *N*-glycans. (Hex5HexNAc4NeuAc1 + 1$SO_4$ + 2Na *m/z* 2056.745; Hex5HexNAc4 + 1$SO_4$ + 2Na *m/z* 1765.19; Hex3HexNAc4 + 1$SO_4$ + 2Na *m/z* 1441.4143; Hex3HexNAc3 + 1Na *m/z* 1136.3964 within a ± 6.0ppm mass error). Serum was spotted on amine reactive slides and separate spots of serum were treated with Neuraminidase A and β-galactosidase to expose GlcNAc residues followed by treatment with a GlcNAc-6-$SO_4$ specific hexosaminidase (F10-ORF19p, Chuzel et al., 2021). *N*-glycans were then released using PNGaseF. Key: *N*-acetyl glucosamine (GlcNAc), mannose (Man), galactose (Gal), sialic acid (Sia), sulfate (S).

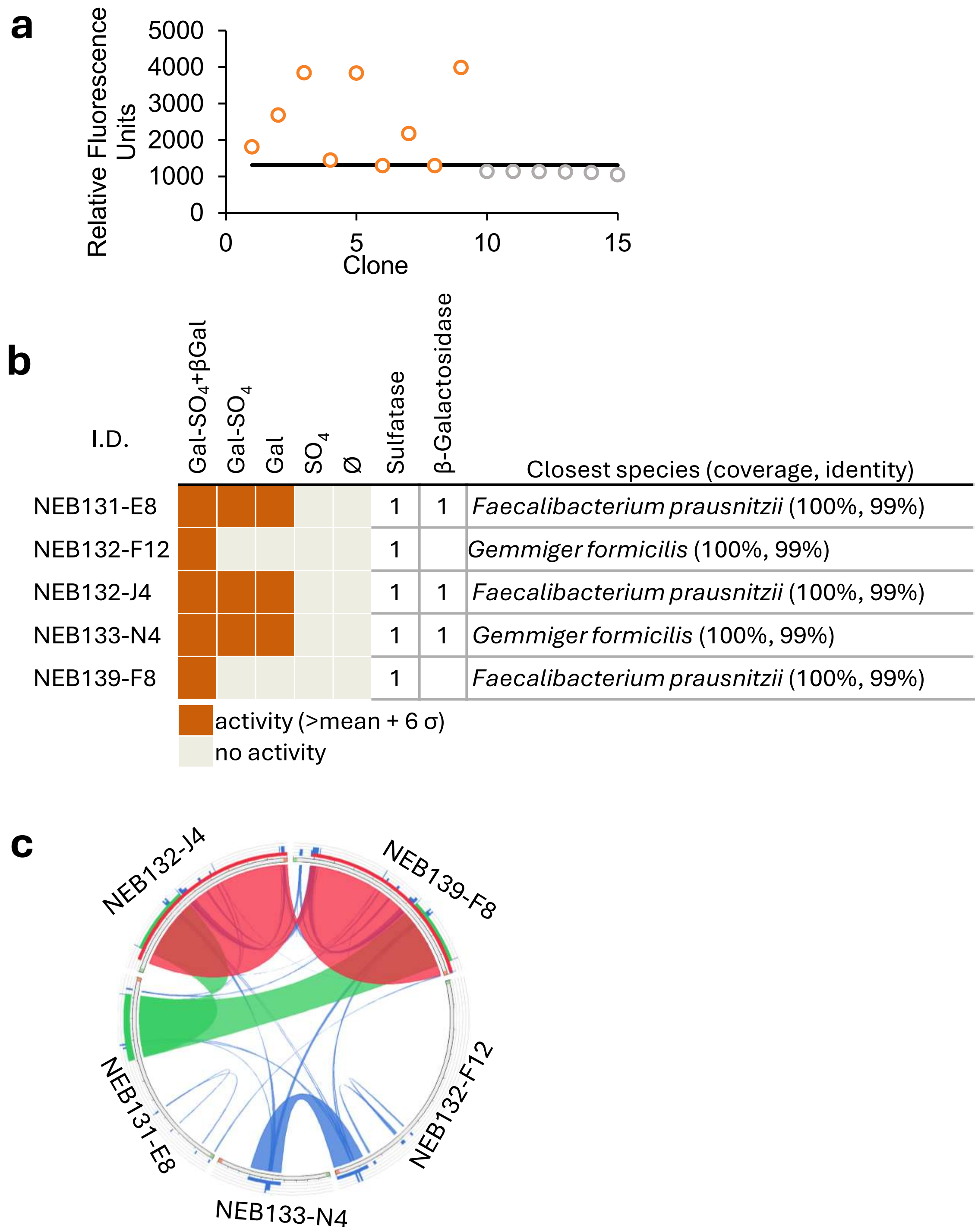


**Supplemental Figure 2. Functional metagenomic screening for sulfatases that act on galactose-3-SO$_4$. a)** Clones of interest from the initial screen (orange) were evaluated for reproducible activity. Replicates of the empty pCC1 fosmid vector (grey) were screened as a control for background fluorescence. Clones exhibiting fluorescence at least 6 standard deviations above the mean background fluorescence (black line) were considered significant. Data shown are the fluorescence following 24h of incubation. **b)** The top 5 clones from the rescreen were tested with four conditions: 4MU-Gal-3-SO$_4$ with exogenous hexosaminidase (Gal-3-SO$_4$+βGal), 4MU-Gal-3-SO$_4$ in the absence of exogenous enzyme (Gal-3-SO$_4$), asulfated 4MU-Gal (Gal), and 4MU-SO$_4$ (SO$_4$). Clones were sequenced and the number of sulfatase and galactosidase genes in each fosmid and the closest species of origin predicted. **c)** A Circos plot made with Circoletto software illustrates relatedness in the fosmid sequences and reveals two unique sulfatases. Overlapping colors indicate regions with identical sequences. The green region, repeated in 3 fosmids, contains 1 unique sulfatase sequence which could be responsible for the activity seen. Likewise, the blue region, repeated in 2 fosmids, contains another unique sulfatase sequence.

a

b

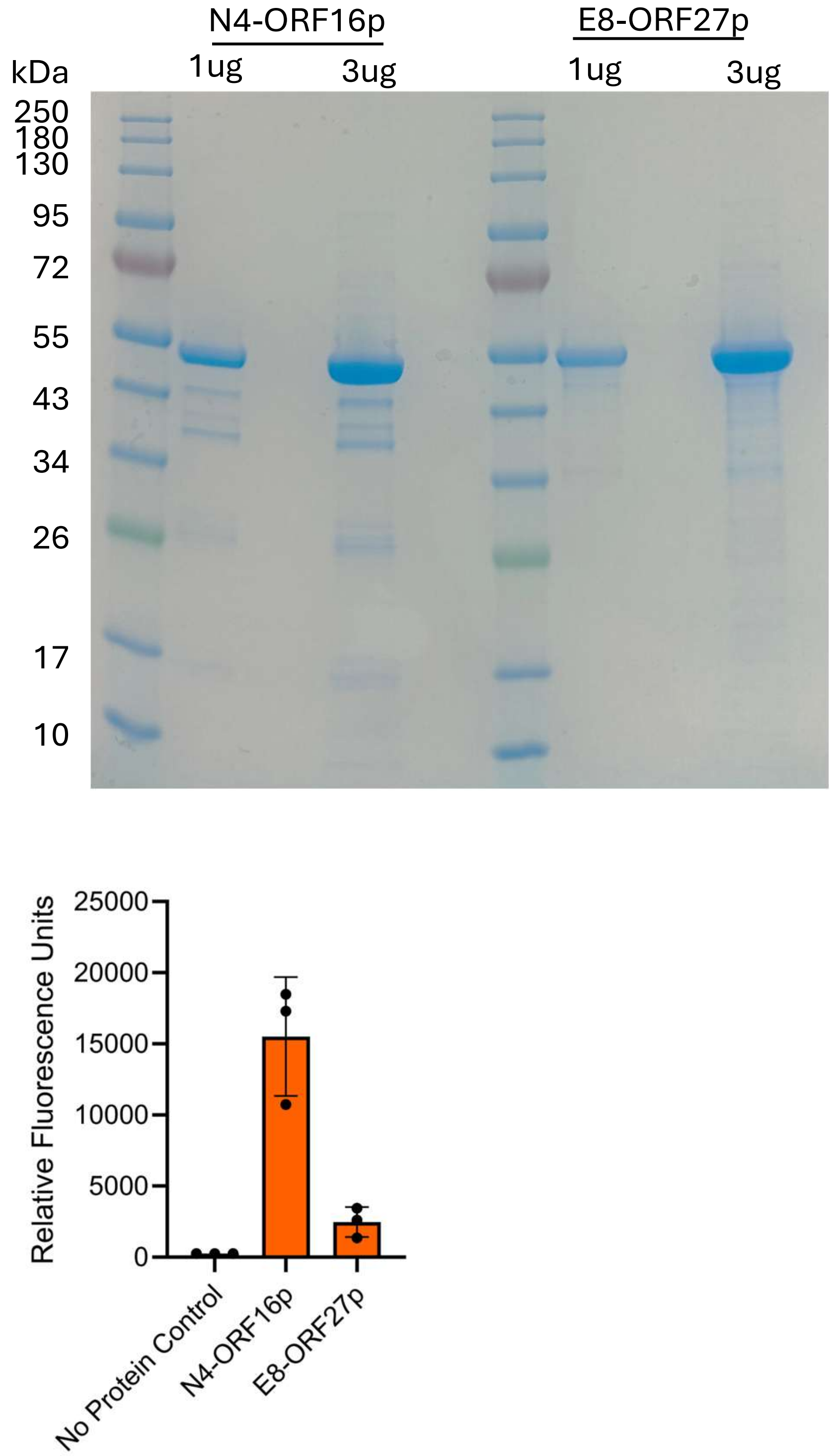


**Supplemental Figure 3. Purified sulfatase candidates. a)** Candidates were expressed with a C-terminal His tag and purified by nickel affinity chromatography. **b)** Activity of purified sulfatase candidates on 4MU-Gal-3-$SO_4$. Assay performed under initial screen conditions, not the optimal reaction conditions later determined, see **Figure S4**. Data presented as mean and standard deviation, N = 3.

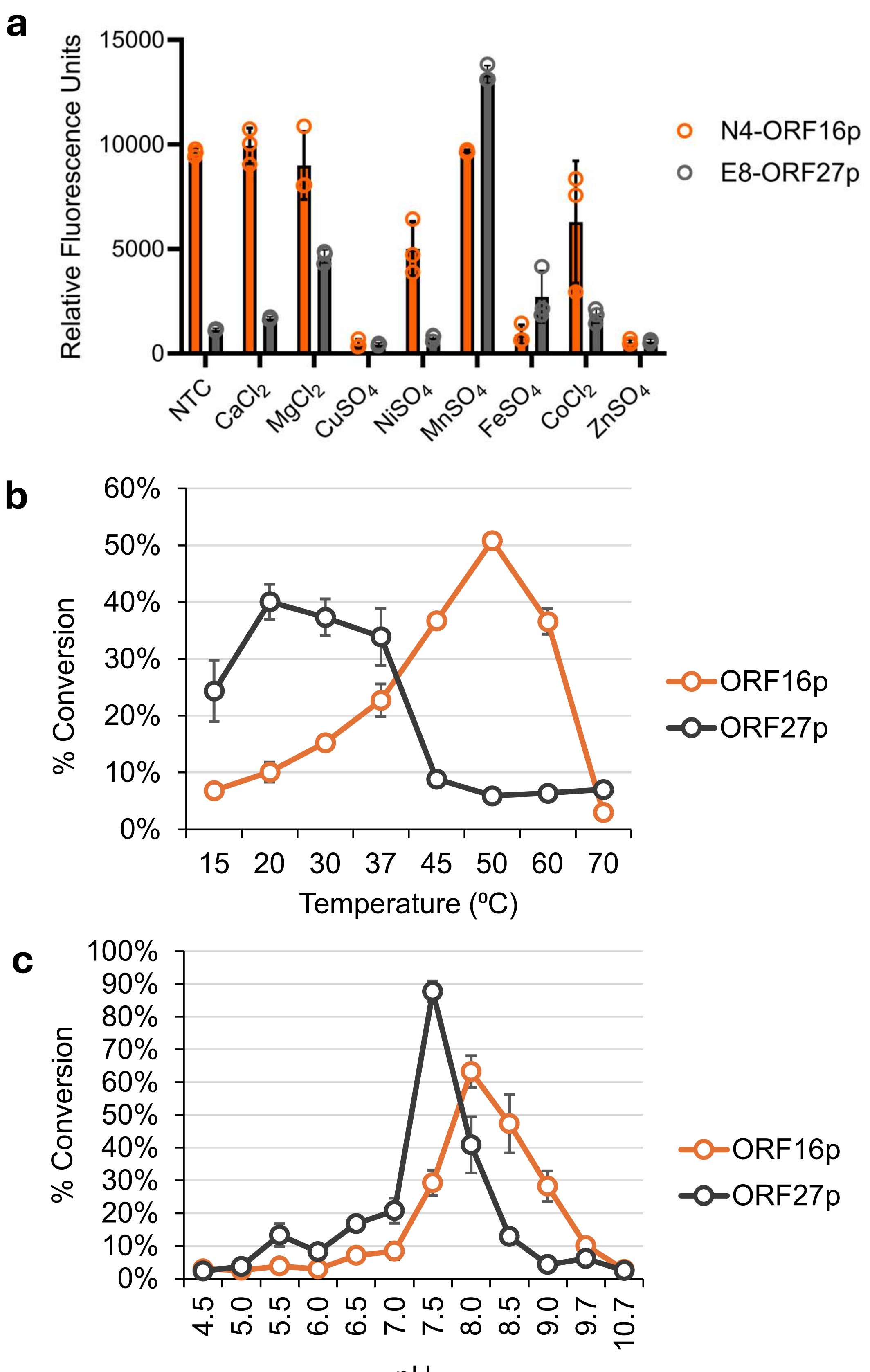


**Supplemental Figure 4. Biochemical characterization of galactose-3-sulfatase candidates.** Purified protein was used to determine optimal reaction conditions for each sulfatase candidate. **a)** The 4MU-Gal-3-$SO_4$ was used to evaluate the effect of metal ions on sulfatase activity. Gal-3-$SO_4$ labeled with procainamide was used to identify the optimal **b)** temperature and **c)** pH, shown as the percent conversion from sulfated to unsulfated. NTC = No Treatment Control, no metal ion added. Data presented as mean and standard deviation, N = 3.

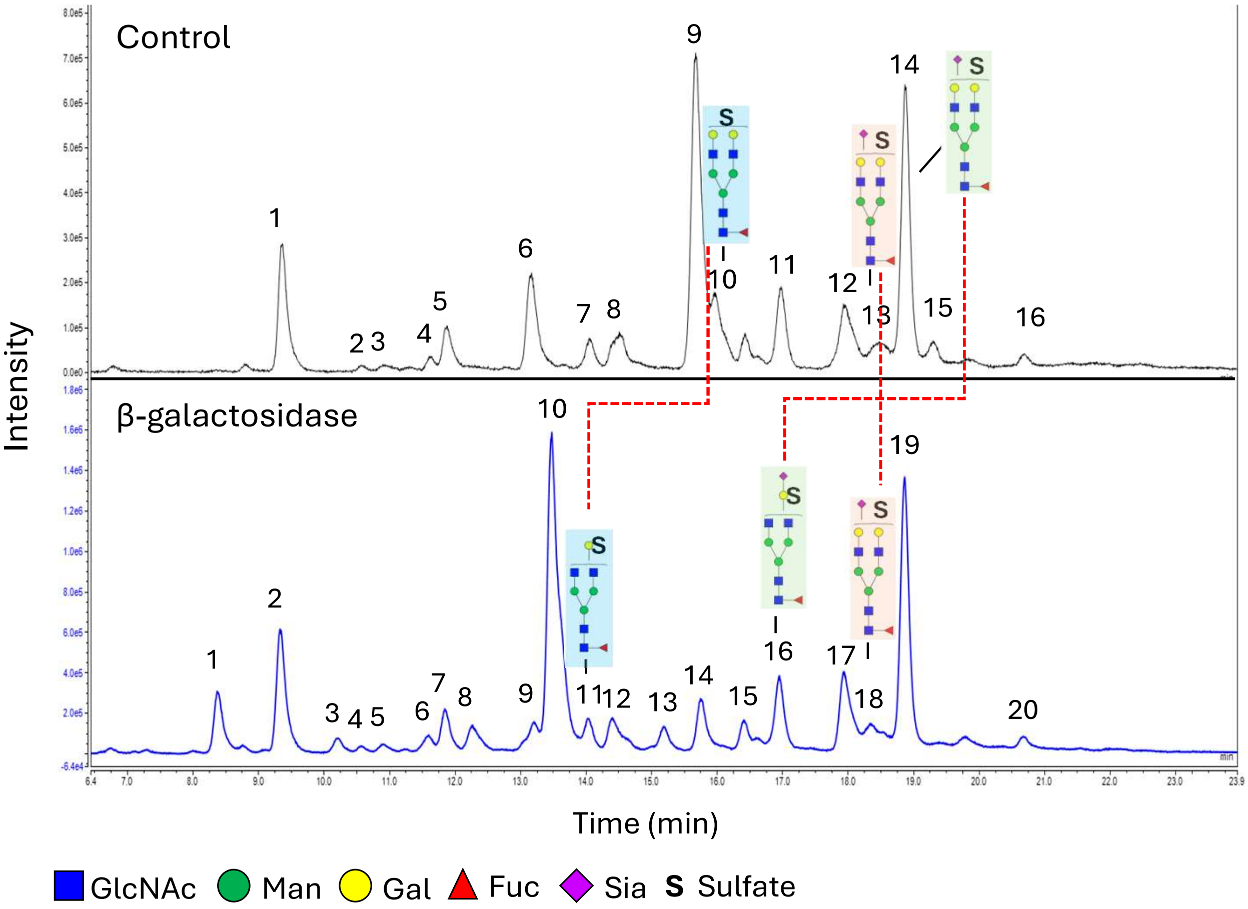


**Supplemental Figure 5. Confirmation of sulfate localization on released thyroglobulin *N*-glycans.** *N*-glycans released from thyroglobulin with PNGase F were labeled with 2-AB, digested with β-galactosidase, and analyzed by LC/MS to confirm localization of sulfates on galactose. *N*-glycan structures were assigned based on previously published thyroglobulin N-glycan structures and detected masses (**Table S4**) (Kayili & Salih, 2022). Key: *N*-acetyl glucosamine (GlcNAc), mannose (Man), galactose (Gal), fucose (Fuc), sialic acid (Sia), sulfate (S).

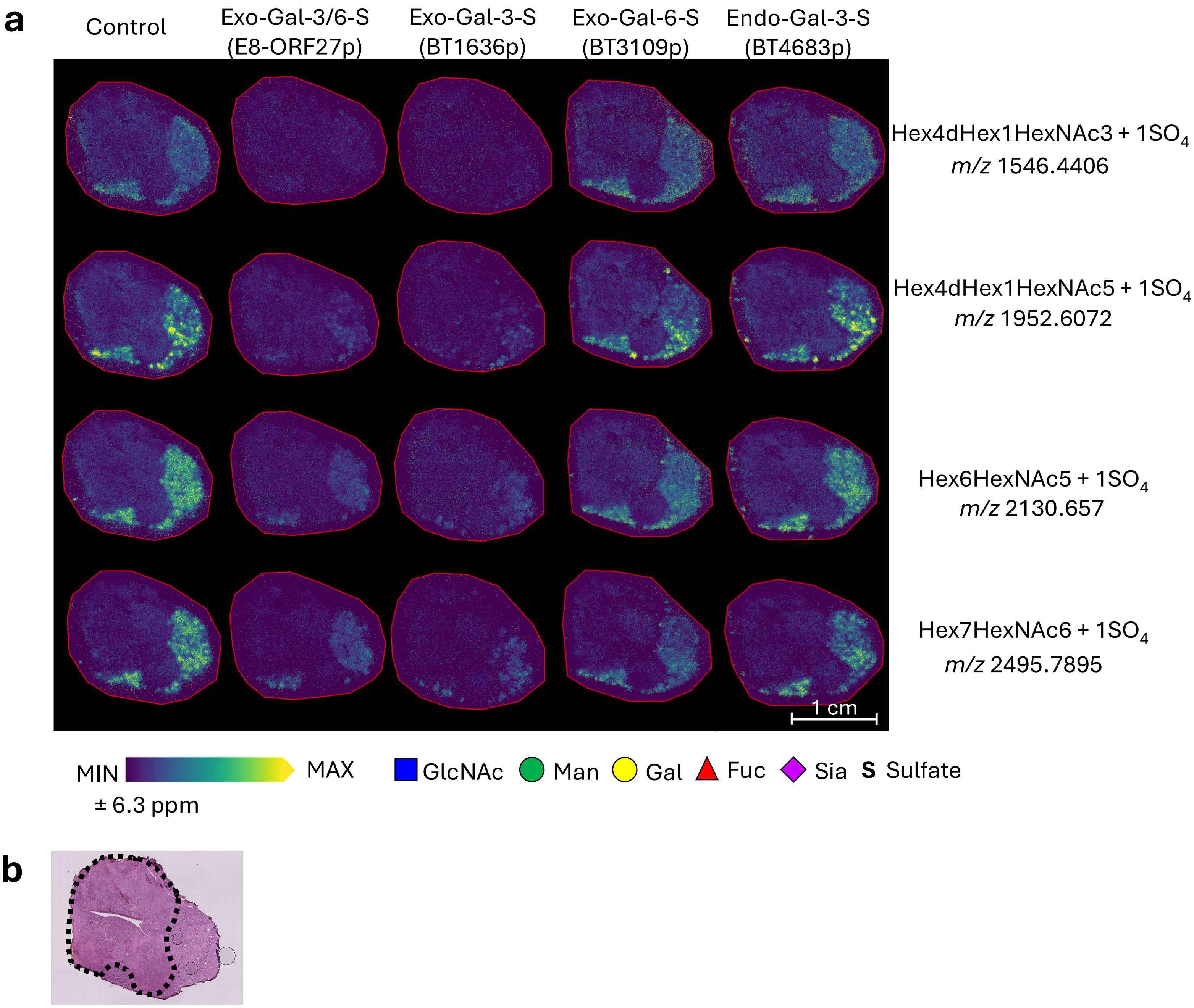


**Supplemental Figure 6. Identification of additional sulfated *N*-glycans identified in cancerous human thyroid tissue. a)** MALDI imaging of sulfated *N*-glycans (Hex4dHex1HexNAc3 + $1SO_4$ + 2Na *m/z* 1546.4406; Hex4dHex1HexNAc5 + $1SO_4$ + 2Na *m/z* 1952.6072; Hex6HexNAc5 + $1SO_4$ + 2Na *m/z* 2130.657; Hex7HexNAc6 + $1SO_4$ + 2Na *m/z* 2495.7895 within a ± 6.3 ppm mass error) present in cancerous human thyroid tissue. Serial sections were treated with either an endo-galactose-3/6-sulfatase (E8-ORF27p), an exo-galactose-3-sulfatase (BT1636p), an exo-galactose-6-sulfatase (BT3109p) or an endo-galactose-3-sulfatase (BT4683p) (Luis et al., 2021) followed by treatment with PNGaseF to release *N*-glycans. **b)** H&E stain of a neighboring section, cancerous region is outlined in black. Key: *N*-acetyl glucosamine (GlcNAc), mannose (Man), galactose (Gal), fucose (Fuc), sialic acid (Sia), sulfate (S).
